# Response plasticity of *Drosophila* olfactory sensory neurons

**DOI:** 10.1101/2021.12.06.471362

**Authors:** Lorena Halty-deLeon, Venkatesh Pal Mahadevan, Bill S. Hansson, Dieter Wicher

## Abstract

In insect olfaction, sensitization refers to the amplification of a weak olfactory signal when the stimulus is repeated within a specific time window. In the vinegar fly, *Drosophila melanogaster*, his occurs already at the periphery, at the level of olfactory sensory neurons (OSNs) located in the antenna. In our study, we investigate whether sensitization is a widespread property in a set of seven types of OSNs, as well as the mechanisms involved. First, we characterize and compare differences in spontaneous activity, response velocity and response dynamics among the selected OSN types. These express different receptors with distinct tuning properties and behavioral relevance. Second, we show that sensitization is not a general property. Among our selected OSNs types, it occurs in those responding to more general food odors, while OSNs involved in very specific detection of highly specific ecological cues like pheromones and warning signals show no sensitization. Moreover, we show that mitochondria play an active role in sensitization by contributing to the increase in intracellular Ca^2+^ upon weak receptor activation. Thus, by using a combination of single sensillum recordings (SSR), calcium imaging and pharmacology, we widen the understanding of how the olfactory signal is processed at the periphery.

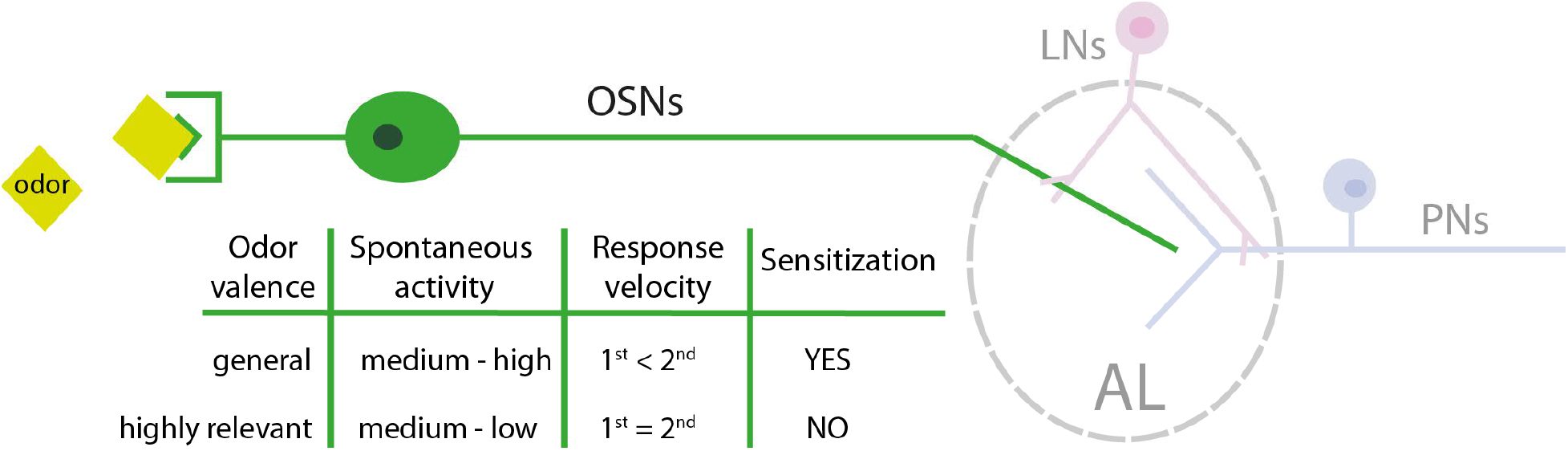

## Introduction

Encoding sensory stimuli is metabolically expensive. Sensory systems have therefore evolved to minimize the cost involved, while maximizing the amount of information gained from a stimulus by adjusting to changes in the environment based on recent input history (Laughlin, 1981). A common feature of sensory systems is the ability to adapt, i.e. to decrease the response to a constant or repeated stimulus. On the other hand, repetitive stimuli can induce sensitization, which leads to a progressive amplification of the response. A recent study states that sensitization leads to a more detailed resolution of the stimulus itself, giving a better representation of the external information (Młynarski and Hermundstad, 2018).

The first report on sensitization was presented in a non-associative learning context on behavioral arousal, in which a novel, strong or noxious stimulus led to an increase in reflex responsiveness (Carew et al., 1971). This phenomenon has been extensively studied in the gill-withdrawal reflex of the sea slug *Aplysia*. In this marine invertebrate, a repeatedly noxious stimulus sensitizes the siphon withdrawal reflex (Pinsker et al., 1973; Hawkins et al., 1998, 2006) through presynaptic facilitation of mechanoreceptive sensory neurons (Castellucci and Kandel, 1976; Klein and Kandel, 1978, 1980).

A form of long-term sensitization has been reported in several organisms during maturation. An increased sensitivity of the olfactory system is associated with reproductive hormone levels in *Drosophila melanogaster*, where older males display higher sensitivity to pheromones (Lin et al., 2016). Exposure to an odorant during a sensitive imprinting period in salmons, resulted in an increased sensitivity to that odorant lasting for months, which was crucial for a successful return to their natal site (Nevitt et al., 1994; Dittman et al., 1997). In another scenario, sensitization can be observed in males of the noctuid moth *Spodoptera littoralis* upon brief pre-exposure to female sex pheromones in behavioral assays (Anderson et al., 2003, 2007; Guerrieri et al., 2012) or upon brief predator sound exposure, a case of cross-modal sensitization (Anton et al., 2011). These experiments indicate that the olfactory system can be modulated by experience-driven plasticity.

Here we focused our interest on a form of olfactory short-term sensitization described in *Drosophila melanogaster*. Sensitization in this case refers to an increased response to weak odor stimuli when repeated within a short time window (Getahun et al., 2013). The olfactory world of insects is highly dynamic. Once emitted from the source, volatiles are dispersed and diluted in the ambient air resulting in a filamentous plume (Murlis et al., 1992). Using a set of ~60 odorant receptors (ORs) (Couto et al., 2005), *Drosophila melanogaster* flies are able to extract valuable information from the plume in terms of odor identity and intensity (Bhandawat et al., 2010a). They are capable of resolving fast changes in odor pulses (Getahun et al., 2012; Szyszka et al., 2014) and can adjust their sensitivity based on previous odor stimuli (Nagel and Wilson, 2011; Getahun et al., 2013). ORs form heteromeric complexes of an odor-specific OR (ORX) and a co-receptor termed Orco (Larsson et al., 2004; Benton, 2006) and are expressed in olfactory sensory neurons (OSNs) housed in the antenna and maxillary palps (Joseph and Carlson, 2015). It has been shown that different OSN types can be sensitized upon repetitive low intensity stimulation (Getahun et al., 2013; Mukunda et al., 2016). However, as OSNs are strongly polarized cells, different signalling mechanisms might exist across the different compartments of the neuron (i.e. soma, inner dendrite, outer dendrite). For example, sensitization at the outer dendrite of the OSN type expressing the Or22a receptor is calmodulin (CaM) dependent, but there is yet at least one more type of regulation in the other compartments that remains elusive (Mukunda et al., 2016). The so far known mechanisms involved in these processes have been reviewed recently (Wicher, 2018). Briefly, an odor stimulus too weak to produce a robust OR activation can lead to cAMP production (Miazzi et al., 2016). This messenger activates Orco, which causes Ca^2+^ influx. Orco activity can then be further enhanced via Ca^2+^-activated calmodulin (CaM) (Mukunda et al., 2014, 2016) and/or via PKC phosphorylation (Sargsyan et al., 2011). These processes finally result in OR sensitization.

Despite the progress made in understating sensitization in olfaction, fundamental questions still remain unanswered. Whether this regulatory process is common for all OSNs or whether it relies on the functional role of an OSN remains unknown. Regarding mechanisms, increase of intracellular Ca^2+^ could also be provided by intracellular stores such as mitochondria, which has been shown to play a crucial role in the olfactory response in both mammals and *Drosophila* (Fluegge et al., 2012; Lucke et al., 2020). However, a possible role of mitochondria in sensitization still remains elusive.

With a combination of single sensillum recordings (SSR), calcium imaging and pharmacology we aim at widening the understanding of how the olfactory signal is processed at the periphery. Here, we first demonstrated differences in the response properties among the studied OSN types and we were able to show that sensitization is not a general property. Among our selected OSN types sensitization was observed among OSNs expressing ORs tuned towards food odors and taking part in cross fibre coding. Furthermore, calcium imaging and pharmacological experiments demonstrated that mitochondria play an active role in sensitization, contributing to the increase in intracellular Ca^2+^.

## Results

### Differential response characteristics between OSN types

As repeated low intensity odor stimulation has been observed to lead to gradually increasing responses, we asked whether this sensitization phenomenon is a general property of OSNs. SSR enables analysis of neuronal activity (Clyne et al., 1997), so we set out to investigate the occurrence of sensitization in a reduced group of OSN types by using this method. SSR data was obtained from a representative set of OSN types responding to food odors (ab3[Or22a, Or85b], ab5[Or82a, Or47a]), pheromone (at1[Or67d]) and danger signals (ab4[Or7a, Or56a]) (**Figure 1A**). We chose these OSN types as representatives not only because of the different behavioral significance of the stimuli they encode (food, danger, mating) but also because the receptors show different tuning properties (narrowly vs broadly tuned). Ab4B, at1 and ab5A represent narrowly tuned receptors, whereas ab3A&B, ab4B and ab5B respond to a wide variety of odors (Hallem and Carlson, 2006). Thus, it should be possible to find out whether such differences are related to the tuning of the receptor itself, to the meaning the odor has for the fly, or to both.

**Figure 1.**
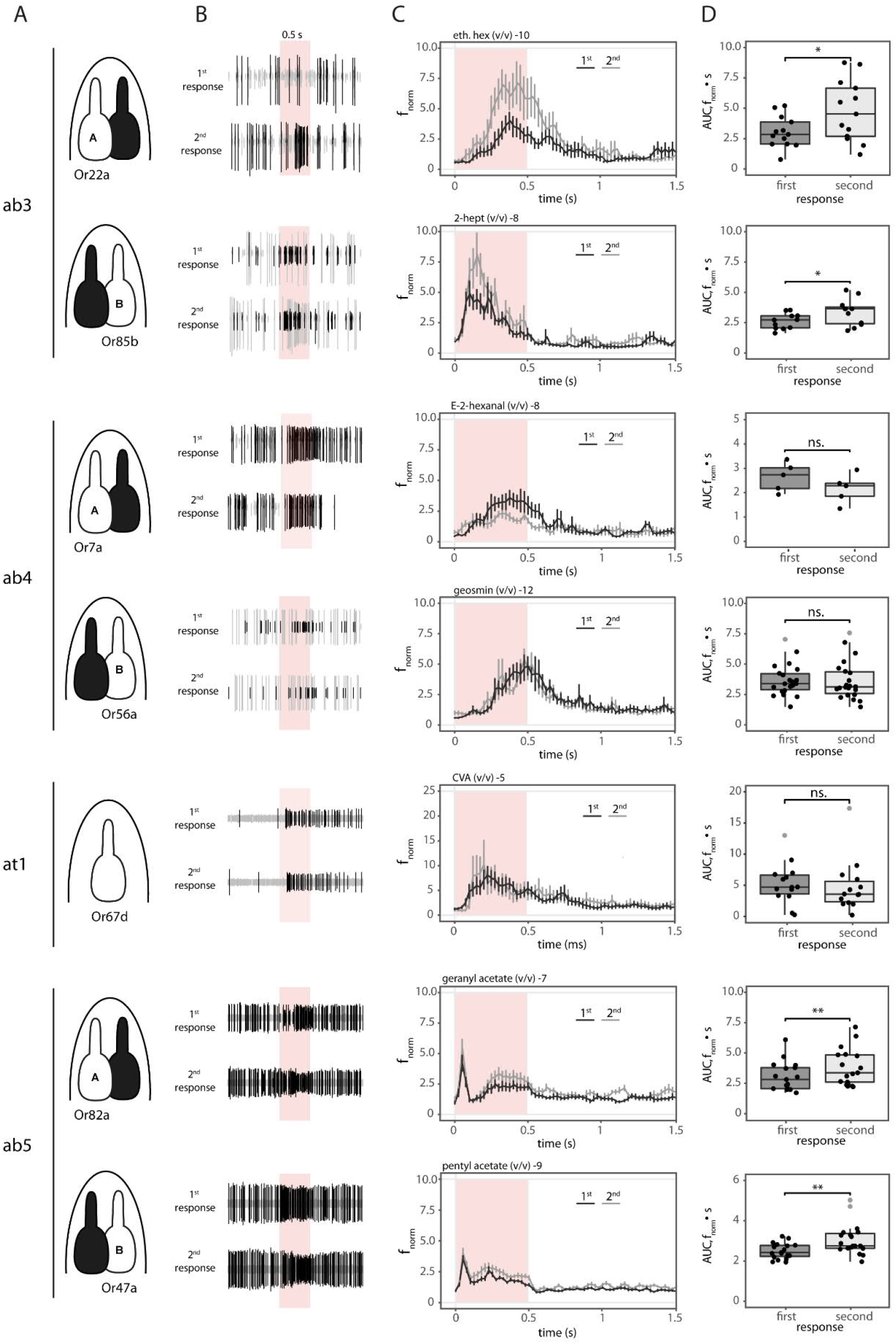
Single Sensillum Recordings. **A** Sensillum of interest (e.g. ab3) and schematic drawing of the neuron that it’s being recording from in white, partner neuron in black. **B** Neuronal activity of the different neurons as in A to two 0,5s stimulation (red bar) 20s apart. **C** Normalized spiking frequency (f_norm_) for first (black) and second (gray) stimulation as in B. Red bar indicates odorant stimulus. Data represent mean ± SEM. **D** Averaged Area Under the Curve (AUC) over 2 seconds corresponding to those in C. Gray dots indicate outliers. Paired t-test: AUC_Or22a n=13 pairs, AUC_Or85b n=11 pairs, AUC_Or7a n=5 pairs, AUC_Or56a n=22 pairs, AUC_Or82a n=17 pairs. Wilcoxon matched-pairs signed rank test: AUC_Or67d n=14 pairs, AUC_Or47a n=19 pairs. *P<0.05, **P<0.01, ns not significant.

We recorded time course, quantified the spontaneous spiking activity and the response velocity, analyzed spiking frequencies and calculated median responses upon odor stimulation with low concentrations of the respective best ligands. All seven neuronal types showed different dynamics (**Figure 1B**). We first quantified the spontaneous activity of all neurons one second before the odor stimulation. It has been previously shown that the spontaneous activity of a neuron is determined by the OR expressed (Hallem et al., 2004; Utashiro et al., 2018), and our results are in agreement with these observations. Spontaneous activity (spikes/s) varied significantly among neurons (Kruskal-Wallis chi-squared =150.38, df = 6, P < 0.0001; **Figure S1**). It is important to note that in the case of ab5, no distinction between ab5A and ab5B was possible due to the very similar spiking amplitude of both neurons. Despite the fact that spikes were considered together, ab5A is narrowly tuned to geranyl acetate, which ensures a differential response to that of ab5B allowing for proper separation of the neuronal spiking activity upon simulation. Partner neurons in ab3 and ab5 showed similar spiking. In contrast, ab4B had a significantly lower spontaneous activity as compared to ab4A. No differences in the spontaneous spiking activity between at1, ab3 and ab4 were found. However, probably due to the fact that spikes were considered together, neurons in ab5 present a significantly higher spiking rate than the others (except ab4A) (**Figure S1**, for detailed statistics see **Table S1**).

To better assess the differences in temporal response pattern a peri-stimulus time histogram (PSTH) was calculated (Olsson et al., 2011). Then, to accurately resolve response kinetics, spiking activity was normalized to 2 seconds before odor stimulation, providing a normalized spiking frequency (f_norm_, **Figure 1C**). This analysis revealed differences in response dynamics. Ab3A, ab4A&B and at1 displayed a slow rising and longer transient phase as compared to ab3B, which showed faster dynamics. In contrast, neurons in ab5 sensilla displayed a fast On and fast OFF response followed by a semi-plateau. Calculation of the response velocity for each OSN type (**Figure S2**) revealed that the second response tended to be faster in ab3A&B and ab5A. To the contrary, for ab4A&B, at1 and ab5B, first and second responses were equally fast.

Calculation of the total area under the response kinetics curve (AUC) for each neuronal type allowed for characterization of median responses between the first and the second odor stimulation (**Figure 1D**). Sensitization is observed if the second response is significantly stronger than the first to the same odor concentration (Getahun et al., 2013). We detected this phenomenon for food-odor-detecting neurons in ab3 and ab5 sensilla (**Figure 1C** and **Table 3**). In contrast, for neurons in ab4 and at1 sensilla (responding to danger signals and pheromones, respectively) we failed to observe any sensitization (**Figure 1C** and **Table 3**). Thus, based on the physiological identity of the neurons studied, we can conclude that for our set of neurons sensitization occured in OSNs responding broadly to food odors irrespective of the OR tuning.

For a more extensive analysis, we set out to compare one sensitizing to one non-sensitizing neuron morphologically and functionally. We chose Or22a as a sensitizing representative since it is the best investigated OR (Dobritsa et al., 2003; Benton et al., 2006; Pelz et al., 2006; Wicher et al., 2008; Aguadé, 2009; Miazzi et al., 2016) and previous studies have confirmed sensitization occurring in these neurons (Getahun et al., 2013; Mukunda et al., 2016). Due to the ecological relevance of geosmin to fruit flies as a highly specific danger signal (Stensmyr et al., 2012) we chose Or56a as the non-sensitizing representative.

### Functional differences between Or56a and Or22a expressing OSNs

To characterize functional differences between Or22a- and Or56a-expressing OSNs we first compared the spontaneous spiking activity from the SSR measurements. Or22a displayed a higher spontaneous activity as compared to Or56a (**Figure 2A**). Then, to better asses OR performance we determined the odor concentration response relationship in calcium imaging experiments. The advantage of imaging compared to SSR is that it allows application of a highly defined concentration of odors. Our concentration dependent results for Or22a-expressing neurons were in line with those described by Pelz et al., 2006 and Jain et al., 2021. In addition we expanded the existing dose response curve for geosmin, where the previously lowest tested concentration was 10^-8^ (Stensmyr et al., 2012), by adding responses down to 10^-10^. Consistently, SSR results and Ca^2+^ imaging dose response experiments showed Or56a-OSNs being 2 order of magnitude more sensitive as compared to Or22a-expressing ones (**Figure 2B**).

**Fig.2.**
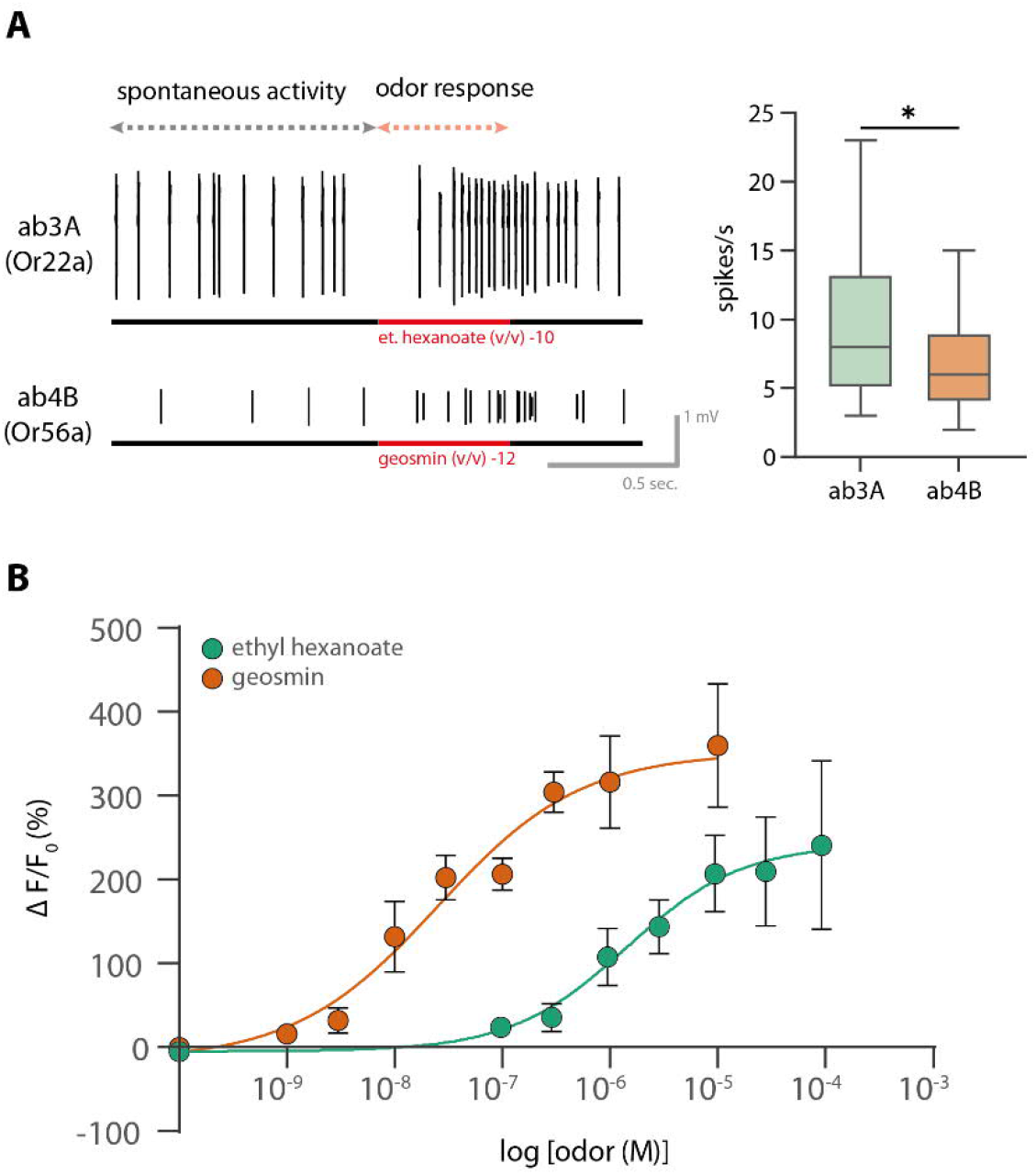
**A Left:** Example of an SSR trace for Or22a-(top) and Or56a-(bottom) expressing neurons in a 2 seconds time window. Spontaneous activity for 1 s is shown before the odor onset (lasting 0.5 s, indicated by the red bar). **Right:** Spontaneous activity for 1 s is significantly lower in ab4B neurons (6.8 ± 0.6 spikes/s) compared to ab3A neurons (9.6 ± 1.1 spikes/s) (Mann-Whitney test, two-tailed, *p < 0.05, n_Or22a_=26, n_Or56a_=44). **B** Ca^2+^ imaging concentration dose response curves for Or22a (green, n_et.hex_=12) and Or56a (orange, n_geos_=9) expressing neurons. Measurements were done in the soma. Curves represent sigmoidal fits described by Hill coefficient 0.8 (eth.hex), 0.6 (geosmin), and EC_50_ of 1.5 μM (et.hex) and 0.025 μM (geosmin). Data represent mean ± SEM.

### Morphological differences between Or56a- and Or22a-expressing OSNs

Recent morphometric studies have highlighted the differences in size and general morphology among OSNs, where the “A” neuron usually is the biggest of the sensillum pair (Hansson et al., 1994; Zhang et al., 2019; Nava Gonzales et al., 2021). Moreover, Nava Gonzalez and coworkers also report that for basiconic sensilla, the large-spike OSNs - for example ab3A - possess an enlarged inner dendrite enriched with mitochondria. Interestingly, Mukunda et al. (2016) proposed intracellular Ca^2+^ sequestration as an additional mechanism contributing to sensitization in inner dendrites and soma of Or22a expressing neurons. In the outer dendrites however, sensitization solely relied on calmodulin (CaM) -dependent processes. In an attempt to link these observations, we set out to determine and compare the morphology of Or56a- and Or22a-expressing OSNs.

Immunostaining allowed visualization of single OSN morphology exposing an enlargement of inner dendrites in O22a-expressing neurons (**Figure 3A**, top left panel, arrowhead). This enlargement was apparently absent in Or56a-expressing neurons (**Figure 3A**, lower panel). In a morphological study by Shanbhag et al. in 2000, and recently shown by Nava Gonzalez et al. (2021), the inner dendritic segment was described to be filled by mitochondria. To explore this possibility, we co-expressed GFP targeted to the mitochondrial matrix (mito::GFP) together with a dendritic marker (DenMark). This allowed for visualization of mitochondria under the control of OSN Or22a- (**Figure 3B**, top) or Or56a-Gal4 drivers (**Figure 2B**, down). The immunostaining of mitochondria confirmed that inner dendrites of Or22a-expressing OSNs show high mitochondrial abundance.

**Figure 3.**
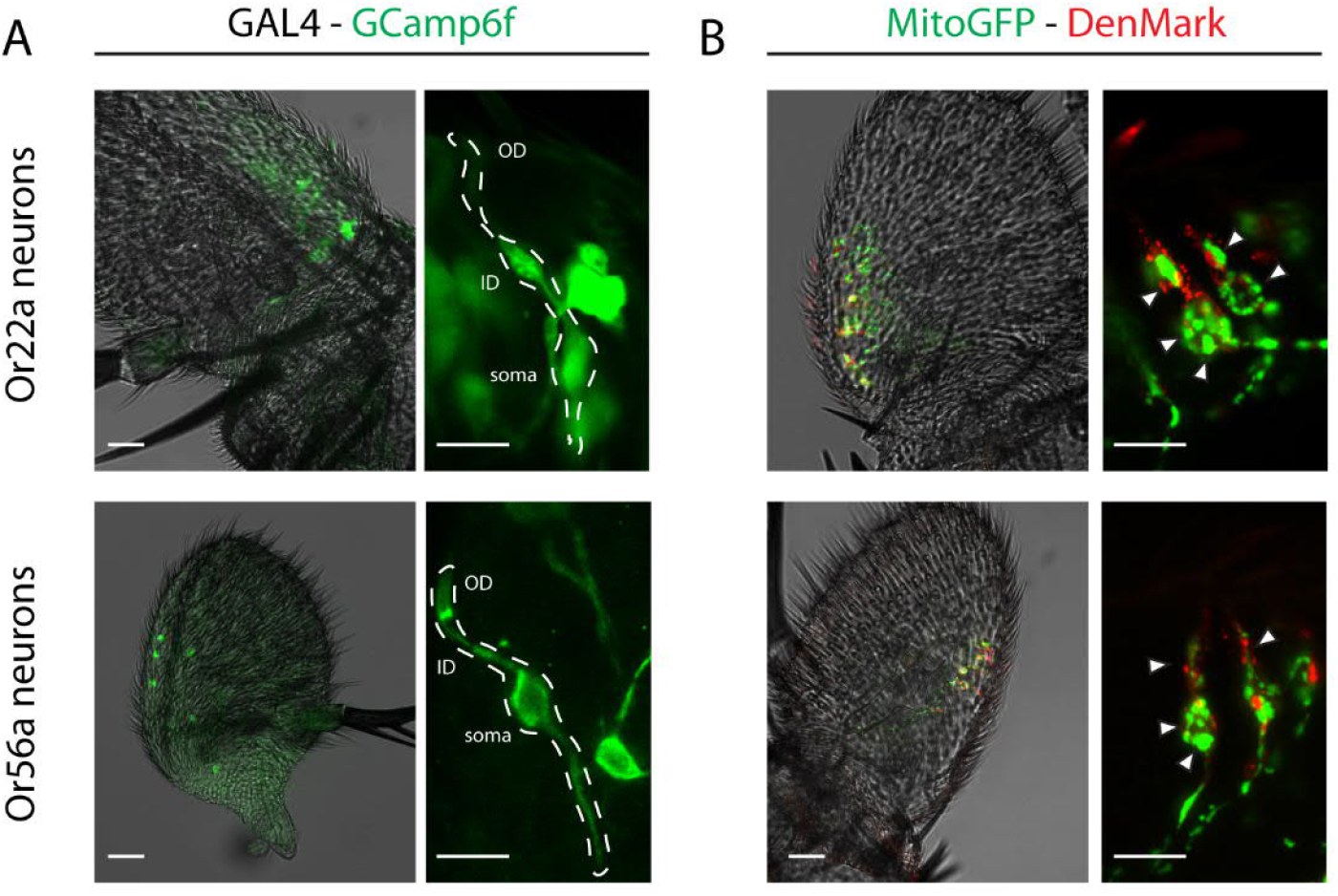
**A** neuronal distribution and morphology of GAL4 lines with the fluorescent marker GCamp6f. Dotted line indicates a single neuron. White arrow indicates inner dendrite. **B** mitochondrial distribution with marked mitochondria (Mito-GFP) and a dendritic marker (DenMark) under the control of the OSN Or22a- (top) or Or56a-Gal4 driver (down). White arrows indicate mitochondria. Scale bar: 20 μm for whole antenna and 10 μm for detail. OD: outer dendrites, ID: inner dendrites.

A quantification analysis of the immunostaining revealed not only that Or22a-expressing neurons have more mitochondria; these OSNs are also larger (**Figure 4**). Inner dendrites or Or22a neurons are significantly enlarged (**Figure 4A**) and heavily packed with mitochondria (**Figure 4B**) as compared to Or56a-expressing neurons.

**Figure 4.**
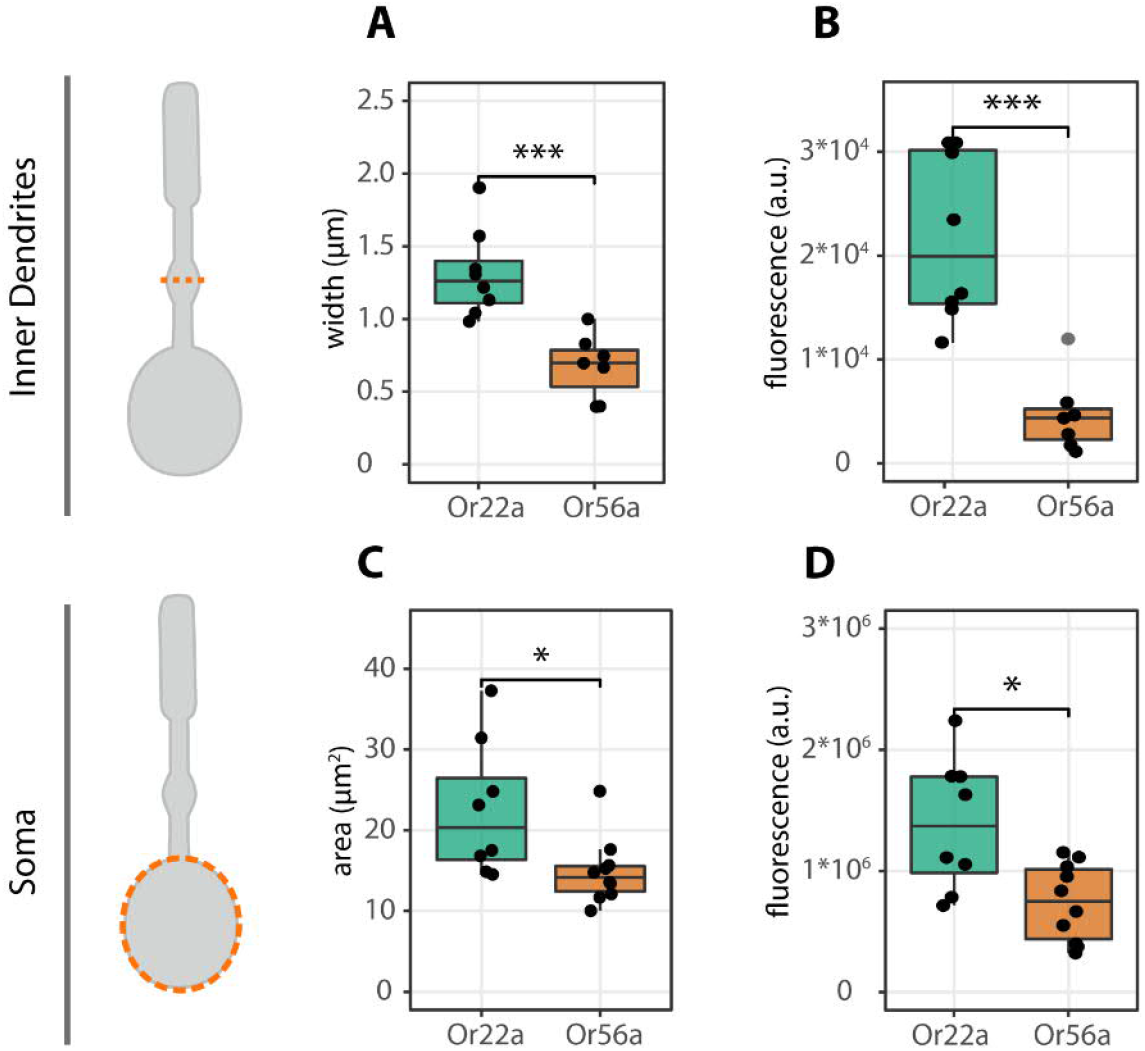
**Left:** schematic drawing of neurons and the compartment analyzed, top panels for inner dendrites and lower panels for somata. Dotted orange line indicates measuring section. **Middle**: width (μm) and area (μm^2^) of inner dendrites (**A**) and soma (**C**) of both neuronal populations. **Left:** corrected total cellular fluorescence (CTCF) calculated as in McCloy et al., 2014 as an estimation of mitochondria abundance. There is a significant difference in the fluorescence intensity between Or22a and Or56a expressing neurons in both the inner dendrites (**B**) and the soma (**D**). Data represent mean ± SEM. Two-tailed t-test, ns, not significant, ***p < 0.001, *p < 0.05. Soma: n_Or22a_=8, n_Or56a_=10; Inner dendrites: n_Or22a_=8, n_Or56a_=7

In the soma, the difference in size is not as pronounced (**Figure 4C**). A difference still exists in the mitochondrial abundance, being more numerous in Or22a expressing neurons also in the soma (**Figure 4D**).

### Sensitization and mitochondria

Following these results, we wanted to evaluate if the observed differences in mitochondria abundance could have an influence in sensitization. The involvement of mitochondria in the *Drosophila* olfactory response has been recently reported (Lucke et al., 2020). Lucke and colleagues found that auranofin, a mitochondrial permeability transition pore (mPTP) activator (Rigobello et al., 2002), caused a significant reduction in the OSN response. In addition, manipulation of mitochondrial function can influence general sensitization of OSNs when stimulated with the Orco agonist VUAA1. The critical player in this case was also shown to be the mPTP, in that application of auranofin depressed sensitization (Wiesel, E. personal communication, November 2021). Therefore, we chose auranofin to evaluate mitochondria influence on sensitization.

Sensitization occurs near threshold and according to the dose responses showed in Figure 2, the chosen concentrations for the experiments were 1 nM (10^-9^) for geosmin and 0.5 μM (10^-7^) for ethyl hexanoate (as in Mukunda et al., 2016). One of the advantages of the open antenna preparation for Ca^2+^ imaging experiments (established by Mukunda and collegues in 2014) is that allows measuring receptor activity in the different compartments of the cell, mainly outer dendrites, inner dendrites and soma. To that end, we designed an experiment consisting of double pared stimulations in an open antenna preparation. One pair stimulation under control conditions and one in the presence of auranofin. This allowed us to compare the direct involvement of mitochondria in the response of our selected cells.

#### Mitochondria are important for sensitization in Or22a-expressing neurons

Sensitization was observed in all compartments of Or22a expressing neurons under control conditions (**Figure 5A-F,** white box; **Table 4**). However, in the presence of auranofin (25 μM), the OR response was diminished and sensitization was abolished (**Figure 5A-F,** pink box; **Table 4**). Furthermore, there was an increase in intracellular calcium [Ca^2+^]_i_ in inner dendrites and soma while in the presence of auranofin, indicating the release of Ca^2+^ from mitochondria (**Figure 5G**). This result indicates that mitochondria play an active role in the sensitization properties of Or22a-expressing neurons.

**Figure 5.**
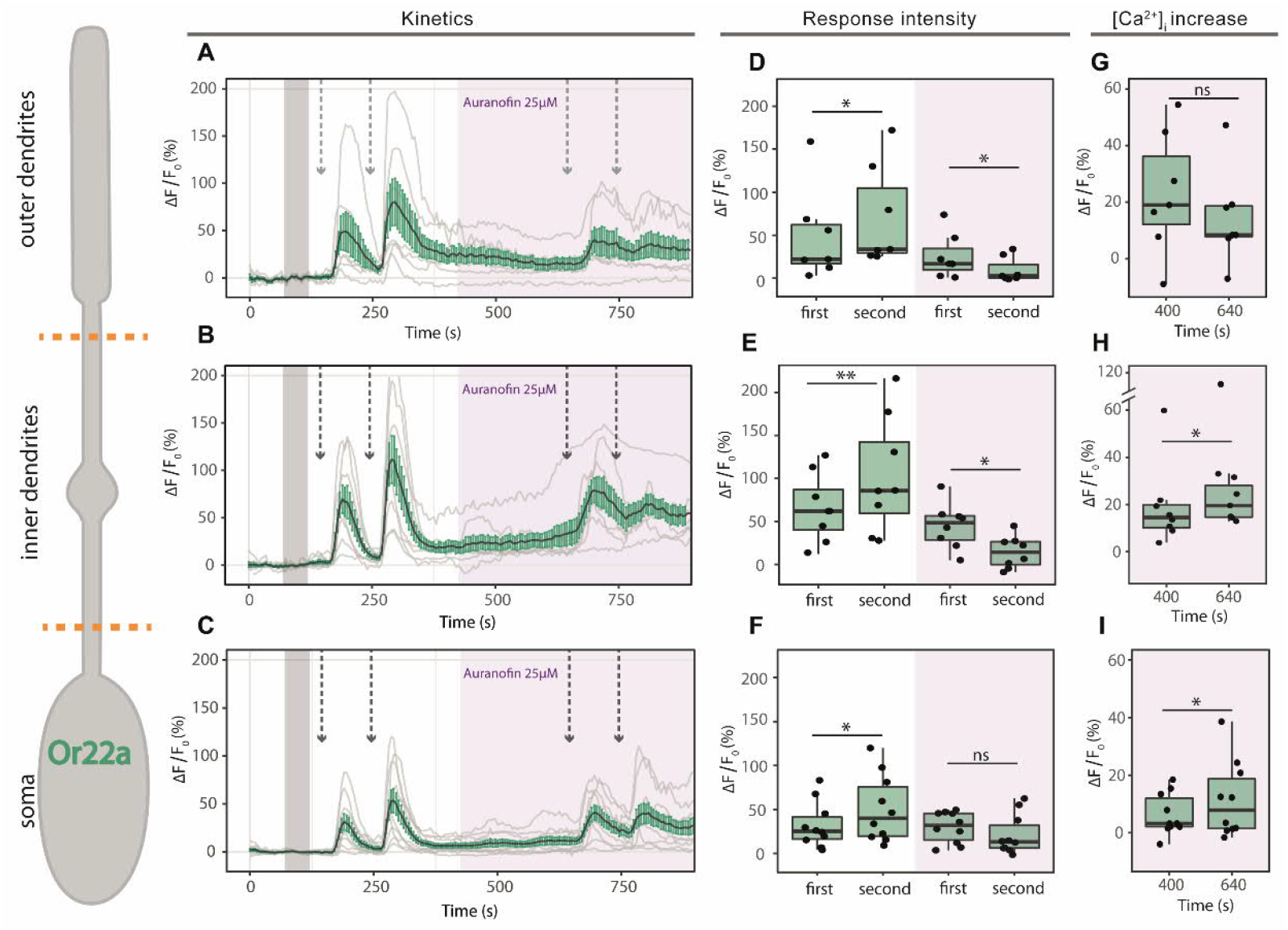
Sensitization in Or22a neurons. Left: schematic drawing of an Or22a-expressing OSNs. Orange dotted lines show division of the different cellular compartments as used for analysis. **A,B,C**: Kinetics show averaged time course of the change in fluorescence intensity (ΔF/F_0_) in *Drosophila* OSNs after application of 0.5 μM ethyl hexanoate (arrows) in outer dendrites (**A**, n=7), inner dendrites (**B**, n=8) and soma (**C**, n=10) under control conditions (in white) and in the presence of the mPTP activator auranofin 25 μM (pink box). Gray bar indicates where data was normalized to obtain ΔF/F_0_. **D,E,F**: maximum increase in ΔF/F_0_ after ethyl hexanoate application in the different compartments as in A-C. **G,H,I**: maximum increase in [Ca2^+^]_i_ between paired stimulations in presence of auranofin 25 μM. Data represent mean ± SEM; one tail paired t-test, ns not significant, * p ≤ 0.05, ** p ≤ 0.01

#### Or56a-expressing neurons show no sensitization

No sensitization event was observed upon stimulation with 1 nM geosmin (**Figure 6A-F** white box, **Table 4**). Then we wondered if sensitization might be visible at even lower concentrations. Hence, we tested gesomin at 0.3 and 0.1 nM in the open antenna preparation. We observed responses only at 0.3 nM and in both cases we failed to observe sensitization (**Figure S2**). Finally, we performed the experiments using gesomin 1 nM, the lowest concentration used for experiments providing strong and reliable responses (**Figure 2B**). In the presence of auranofin (25 μM), the second response was significantly lower only in the soma (**Figure 6A-F** pink box, **Table 4**). Between the 2 pairs of stimulations, there was no change in in [Ca^2+^]_i_ (**Figure 6G**). These results indicate that mitochondria are important in restoring basal calcium levels after stimulation to ensure a proper second stimulation, but play no apparent important role in controlling neither the response intensity nor the intracellular calcium levels.

**Figure 6.**
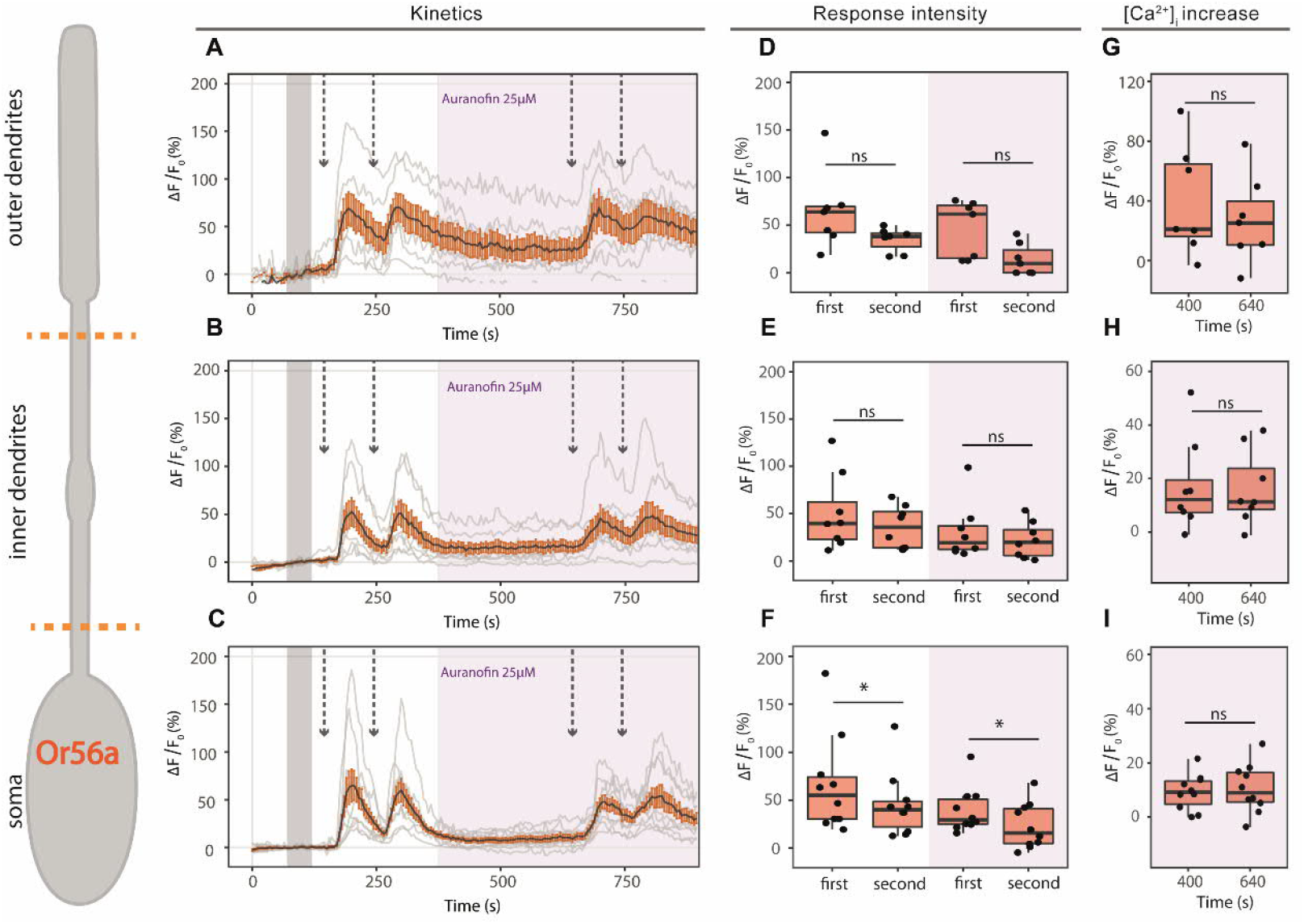
No sensitization in Or56a neurons. Left: schematic drawing of an Or56a-expressing OSNs. Orange dotted lines show division of the different cellular compartments as used for analysis. **A,B,C**: Kinetics show averaged time course of the change in fluorescence intensity (ΔF/F_0_) in *Drosophila* OSNs after application of 0.1 nM geosmin (arrows) in outer dendrites (**A**, n=8), inner dendrites (**B**, n=7) and soma (**C**, n=10) under control conditions (in white) and in the presence of the mPTP activator auranofin 25 μM (pink box). Gray bar indicates where data was normalized to obtain ΔF/F_0_. **D,E,F**: maximum increase in ΔF/F_0_ after geosmin application in the different compartments as in A-C. **G,H,I**: maximum increase in [Ca2^+^]_i_ between paired stimulations in presence of auranofin 25 μM. Data represent mean ± SEM; one tail paired t-test, ns not significant, * p ≤ 0.05, ** p ≤ 0.01.

## Discussion

Understanding how odors are detected and processed at the olfactory periphery is crucial to comprehend how information is then modulated. Here we investigated whether sensitization, an amplification of the olfactory response at the OSN level in the *Drosophila* antenna, is a widespread property in a set of OSNs of the fruit fly.

Remarkably, we found that OSN types expressing different ORs respond to odor stimuli with different strength and dynamics. This indicates that the tuning OrX protein not only determines the resting activity of OSNs (Hallem et al., 2004) but also the characteristics of the odor response. First, and in agreement with others (Hallem et al., 2004), we observed differences in the spontaneous firing of the different OSN types (**Figure S1**). Spontaneous activity originates in the OSNs (Joseph et al., 2012; Stengl and Funk, 2013) where ORs and Orco must be functional, and sensillar components must be intact to generate a baseline firing (Benton et al., 2007). Interestingly, OSNs expressing the most narrowly tuned ORs (Or56a and Or67d) showed the lowest spontaneous activity (**Figure S1**). Sparse code (few spikes) usually allows for a better separation of sensory inputs, however at the expense of being more sensitive to noise (Zhang et al., 2013). *Drosophila’s* OSNs may have found the solution to this problem in that neurons with low spontaneous activity are narrowly tuned to one or few compounds. Thus, sensitivity is increased but the influence of random activation (noise) is diminished. This seems to be the case for pheromone sensing neurons in flies and moths (Kalinová et al., 2001; Dolzer et al., 2003; Benton et al., 2007; Jeanne and Wilson, 2015; Nolte et al., 2016), as well as for highly relevant danger signals for flies (this study).

Second, we observed and quantified differences in response dynamics. (**Figure 1C** and **Figure S2**). As expected for responses near threshold, neurons in ab3, ab4 and at1 (with the exception of ab3B) showed a more tonic response (Martelli et al., 2013). In contrast, responses of neurons in ab5 sensilla displayed a response similar to that of retinal bipolar-ON cells when adapting to light (Masu et al., 1995 their Figure 4 A). These differences, observed in the response kinetics (**Figure 1**), are in agreement with previous studies stating that response dynamics depend on odorant type (Martelli et al., 2013), receptor type (Getahun et al., 2012) and neuron identity (Nagel and Wilson, 2011). The delayed odor-response observed for ab4B and ab3A could be related to the time needed for the odor concentration to reach the neuron’s detection threshold. In the case of geosmin, with a very low vapor pressure (0.001 mmHg v, Stensmyr et al., 2012), it might be that only few molecules reach the OSNs at a concentration of 10^-12^. This highlights the extreme sensitivity of these neurons as already reported elsewhere (Stensmyr et al., 2012). However, to our knowledge, this is the first time such a low concentration has been used for electrophysiological recordings. Although not significant, differences in the response velocity upon two subsequent odor stimulations can be observed (**Figure S2**). For ab4B, at1 and ab5B, first and second responses are equally fast. However, for ab4A, the second response tends to be slightly slower. In contrast, for ab3A&B and ab5A second responses tend to be faster as compared to the first. This duality in response velocity reflects the double odor transduction strategy proposed by Wicher et al. (2008, 2010): a combination of a slower, more sensitive metabotropic with a fast, purely ionotropic pathway.

The described mechanism for sensitization reviewed recently (Wicher, 2018; Wicher and Miazzi, 2021) further supports this view. Upon a first stimulus of low odor concentration, there is a slower and more sensitive metabotropic response in which autoregulative processes through PKC (Sargsyan et al., 2011; Getahun et al., 2016) and cAMP (Miazzi et al., 2016) tune the OR to its deserved sensitivity. As a result, a second stimulation of the same odor concentration causes faster and larger ion fluxes. This has been proven true for a few OSNs (Getahun et al., 2013), and we have investigated whether this phenomenon is a widespread property in our panel of 7 OSNs with different valences and different OR tuning properties.

In our sample, ab4A and ab4B represent OSNs encoding odors with negative valence and at1 for pheromone signals. Remarkably, all three failed to sensitize (**Figure 1** ab4A&B and at1). Or56a-expressing neurons (ab4A) only respond to geosmin, which is produced by microorganisms such as mold fungi (La Guerche et al. 2006) and bacteria (Gerber and Lechevalier, 1965) and has a great ecological relevance for the fly (Stensmyr et al., 2012). To date, geosmin is the only aversive odor found with a dedicated pathway, the other aversive odors are represented by combinatorial activation of glomeruli (Knaden et al., 2012; Stensmyr et al., 2012; Seki et al., 2017). Its partner neuron (ab4A-Or7a), responds best to E2-hexanal, and although it has been classified as partially attractive (Hallem and Carlson, 2006), artificial activation of its target glomerulus DL5 leads to aversive behavior. Therefore Or7a-expressing neurons can also be defined as an aversive input channel (Mohamed et al., 2019). Interestingly, it has been proposed that aversive odors might be processed with a different logic than attractive ones in higher brain centers (Gao et al., 2015). In walking flies, detection of an odor increases the frequency of turning. After turns to aversive odors flies moved more quickly and followed straighter paths away from the source as compared for turns following an attractive odor (Gao et al., 2013). Such “runaway” behavior could shorten the exposure to harm. Therefore, the aversive response may rely on specific activity patterns of individual OSNs (Gao et al., 2015). In line with this, larval OSNs carrying highly relevant information for survival are regulated differently in the AL via local interneurons (LNs). Reduced presynaptic inhibition of OSNs responding to odors associated with life-threatening situations allows *Drosophila* larvae to detect these odors less dependently of the response intensity of other OSNs (Berck et al., 2016). Our results are in line with these observations, where detection of aversive odors at low concentrations is sufficient to elicit a robust OSN response. Once the potential harmful situation is faithfully detected, it is likely that the source will not be further investigated, and an amplification of the signal would consequently no longer be necessary.

Along with geosmin, cVA detection is another well-established example of a dedicated circuitry. Several factors contribute to a high sensitivity of this pathway. First, it is detected solely by Or67d-expressing neurons in at1 sensilla targeting the DA1 glomerulus (Tal and Smith, 2006). These neurons have a low detection threshold thanks to a reduced spontaneous firing activity (Jeanne and Wilson 2015 and Figure S1 of this study). Second, the low detection threshold in combination with a high number of cVA responding neurons (~ 40), renders this neuronal population highly sensitive. Third, the OSNs’ postsynaptic partners in the AL (the projection neurons (PNs)) fire a spike after only a small percentage of the OSNs have responded to a stimulus and are capable of up to 3-fold amplification (Jeanne and Wilson, 2015). As a result, PNs are able to respond rapidly to changes in the number of spikes from the OSNs (Bhandawat et al., 2007; Jeanne and Wilson, 2015). These results, together with the fact that cVA is detected at close range, makes this neural population less susceptible to sensitization. Our results are consistent with this assumption in that, with a concentration of 0.001%, we observed clear responses but failed to see sensitization (**Figure 1** at1).

In contrast, sensitization was observed in OSNs tuned to food odors (**Figure 1** ab3A&B and ab5A&B). Food odors provide crucial information about potential foraging sites, where behaviors such as mating and oviposition occur (Couto et al., 2005; Hallem and Carlson, 2006). Or22a (ab3A) and Or85b (ab3B) represent broadly tuned receptors, responding to many odors. On the other hand, Or82a (ab5A) is narrowly tuned to few compounds, while its partner neuron ab5B houses a broadly tuned receptor (Or47a). The four OSNs project their axons into glomeruli with positive valence in the AL (Or22a➔DM2, Or85b➔VM5d, Or82a➔VA6, Or47a➔DM3) (Knaden et al., 2012). The fact that sensitization occurs in all four OSNs indicates that this property is rather linked to the behavioral significance than to the tuning properties of the receptor. In line with this, a previous study showed that flight responses to odors eliciting attraction are dependent on the identity of the OSNs being activated (Bhandawat et al., 2010b). Bhandawat and colleagues showed that activation of one single neuron type was sufficient to initiate a flight surge even at low concentrations (Bhandawat et al., 2010a).

Behavioral responses to odors during flight are fast and are observed within 100 ms after onset of OSN activity (Bhandawat et al., 2010a). However, Getahun et al. (2013) found that OSN sensitization required an interstimulus interval of 10 s. It has been hypothesized that sensitization could aid flies following faint odor plumes when on a flying search (Getahun et al., 2013, 2016). So how can we reconcile these differences in time domains? Two processes might be happening in parallel, at the OSNs and at the PNs.

OSNs might be subjected to a readiness or awareness state as described by Angioy et. al., (2003). Angioy and colleagues monitored the cardiac activity of moths to evaluate olfactory detection at threshold levels. The heart response accurately indicated odor detection, but an extremely low concentration not suitable for behavioral testing. They postulate that this extreme sensitivity might be due to the formation of awareness to a certain stimulus and the readiness to respond behaviorally. We believe that sensitization follows the same principle. A first reduced response of OSNs puts the system on guard, where a weak odor stimulus leads to activation of the co-receptor Orco by PKC dependent phosphorylation through cAMP (Getahun et al., 2013, 2016; Miazzi et al., 2016). This results in influx of Ca^2+^, which may activate PKC and CaM signaling loops amplifying the Ca^2+^ influx further and thus increasing the sensitivity of the system. As a result, a stronger response upon the same stimuli is observed after the second application.

In parallel, weak OSN inputs are amplified in the PN layer as these neurons respond strongly to small increases in OSN firing rate (Bhandawat et al., 2007). PNs collect information from converging OSNs input in the AL and carry the information to higher brain center such as mushroom body and lateral horn for final processing (Seki et al., 2017). Since PNs pool information from all OSNs expressing the same receptor in its cognate glomerulus, low odor stimuli are detected more quickly and accurately based on a single PN spike response (Bhandawat et al., 2007; Wilson, 2013; Jeanne and Wilson, 2015). This fast encoding mechanism allows the animal to detect the odor onset at a very early phase (Kim et al., 2015). In addition, as with other forms of sensitization (Castellucci and Kandel, 1976; Klein and Kandel, 1978; Appleby and Manookin, 2019), presynaptic modulation can further tune the signal transfer from OSNs to PNs (Wang, 2011; Mcgann, 2013). GABA-dependent presynaptic inhibition has been reported to affect gain control of OSNs through lateral interneurons (LNs). Interestingly, OSNs responding to CO_2_, an innate aversive cue, shows low levels of GABA receptors, which indicates reduced presynaptic inhibition (Root et al., 2008). This may allow for a more fine detection, as seen for *Drosophila* larva (Berck et al., 2016). It would be interesting to evaluate whether sensitizing neurons express higher levels of GABA receptors, indicating higher susceptibility to presynaptic inhibition and lateral modulation.

As a conclusion, and in agreement with others (Gao et al., 2015), we propose that odors of extreme ecological significance, as pheromones and alarm signals, are perceived differently. For these dedicated pathways, there is an investment in more accurate detection at the OSN level; therefore, further sensitization is not needed. However, for food odors, modulation at the OSN level together with the high sensitivity of the PNs to OSN output ensures faithful coding even at low odor concentrations.

This provides a theoretical framework of the use of sensitization for a flying insect. But what are the mechanisms that make sensitization possible in a subset of OSN population? To answer this question we focused on one representative example of each class: Or22a- and Or56a-expressing neurons.

### Mechanisms in sensitization

Getahun et al (2013) postulated that sensitization is intrinsic to the particular OSN type and our results are in agreement with this observation. When we examined sensitization in their native environment, only Or22a-but not Or56a-expressing neurons were sensitized. However, when heterologously expressed in HEK cells, both Or22a and Or56a showed sensitization (Mukunda et al., 2016). In addition, the study of Mukunda et al also showed that sensitization was differently regulated in the distinct compartments of Or22a-expressing neurons, being exclusively CaM-dependent only in the outer dendrites (Mukunda et al., 2016). These results indicate that the receptor alone can be sensitized, but the intrinsic properties of the neuron define the final response. Considerable effort has been spent to understand the diversity of responses of OSNs as a function of their expression of different odor receptors (Getahun et al., 2012; Kolesov et al., 2021). Nevertheless, it has become clear that receptors alone cannot explain the sensitivity, specificity and temporal precision observed in odor-evoked neuronal activity (Slankster et al., 2019; Schmidt and Benton, 2020). OSNs are classified according to their presence in different sensillum types: basiconic, trichoid and coeloconic (Shanbhag et al., 1999) and their responses to different chemical classes (Hallem and Carlson, 2006). Only recently, the morphological features of different OSN types have been systematically examined (Nava Gonzales et al., 2021). This study not only reveals a difference in size between partner OSNs within a single sensillum, but also different dendritic branching and particularly interesting, in mitochondrial abundance. These different morphological aspects of OSNs will likely influence olfactory function.

Our immunostaining results are in agreement with the morphological differences observed by others before (Zhang et al., 2019; Nava Gonzales et al., 2021). Or22a-expressing neurons are bigger and have an enlarged inner dendrite packed with mitochondria as compared to Or56a-expressing neurons (**Figures 3 and 4**). Moreover, in agreement with our hypothesis for higher sensitivity in highly specific OSNs (previous section), geosmin-detecting neurons showed to be two orders of magnitude more sensitive than those detecting ethyl hexanoate (**Figure 2**). These results allowed us to design an experiment in which we could test the influence of mitochondria on sensitization in a selected neuron.

Under control conditions, results from Ca^2+^ imaging experiments are consistent with SSR data, as sensitization is only observed in Or22a-expressing neurons (**Figure 5** and **6**). The general reduction of response intensity observed in the presence of auranofin for both neuronal populations could be due to the presence of Ca^2+^ hotspots in the vicinity of ORs. Ca^2+^ accumulation near the plasma membrane as a result of activation of the mitochondrial permeability transition pore (mPTP) could lead to an early channel closure as known for other Ca^2+^ carrying ion channels (Morad and Soldatov, 2005). Alternatively, reduction in the response could be related to other auranofin effects (Froscio et al., 1989). However, for Or22a-expressing neurons an increase of the intracellular calcium concentration ([Ca^2+^]_i_) indicates auranofin-dependent activation of mPTP. In addition, as a result of auranofin application, sensitization is no longer observed (**Figure 5**). In *Drosophila melanogaster*, mitochondria play an active role as regulators of the odorant response in OSNs (Lucke et al., 2020). Furthermore, mitochondria were shown to be fundamental in shaping OSN response profiles to odors and also in maintaining sensitivity in the olfactory signal process of mammals (Fluegge et al., 2012). We propose that, for Or22a-expressing neurons, Ca^2+^ release from mitochondria could contribute to further activation of Orco through feedback loops mediated by PKC or calmodulin (CaM) (Wicher, 2018). This would drive the OR to a sensitized state resulting in an amplification of subsequent responses. However, upon auranofin-dependent activation of the mPTP, a slow increase in the [Ca^2+^]_i_ occurs. This indicates Ca^2+^ release, as observed before by Lucke et al., (2020). Thus, as Ca^2+^ is no longer stored in mitochondria, there is no contribution of mitochondrial calcium to the response elicited by ethyl hexanoate stimulation and sensitization is no longer present.

In contrast, in Or56a-expressing neurons mitochondrial calcium appears to play a different role. Upon auranofin addition, there was no significant increase in the intracellular calcium concentration **(Figure 6 G,H,I**). We believe that in this case, mitochondria serve mainly as a calcium clearance organelle to ensure a rapid return to basal levels. Influence of mitochondria in sensory response recovery has been previously reported in *D. melanogaster* OSNs (Lucke et al., 2020) and in the photoreceptors of zebrafishes. In cone photoreceptors of zebrafish, mitochondria are tightly clustered between the outer segment and the cell body (Giarmarco et al., 2017). This disposition allows for mitochondrial Ca^2+^ clearance from the outer segment upon stimulation. This process is essential to promote response recovery (Hutto et al., 2020). Similarly, mitochondria present in the soma and inner dendrites serve a function for Ca^2+^ uptake after a first response to ensure a rapid return to basal levels (**Figure 6 B,C, white box**). However, in the presence of auranofin this is no longer possible, thereby resulting in a reduction of the second response (**Figure 6 E,F**).

## Concluding remarks

Evidence that OSNs have a greater role than previously considered in the processing of olfactory signals is growing (Fleischer et al., 2018; Schmidt and Benton, 2020). We expand this knowledge by showing that the first modulation of the olfactory response really occurs at the periphery. In such modulation, behaviorally highly relevant odor information, serving specific purposes such as detection of pheromone or danger signals, are processed differently. However, whether the differences observed at the OSN level are still apparent in the AL, or whether a “signal normalization” (maybe through LNs) ensuring a consistent output occurs is a very interesting question that remains to be investigated.

Furthermore, OSNs of *Drosophila* rely primarily on two types of olfactory receptors, odorant receptors (ORs) and ionotropic receptors (IRs). In the present study we focused on ORs. However, recent investigations have shown that IRs colocalize more widely with ORs than previously thought (Task et al., 2020; Younger et al., 2020). Whether IRs influence the OR response was outside the scope of our study, nevertheless it is an interesting possibility that prompts further investigation. In addition, differences in receptor sensitivity observed throughout this study could be due to the dwelling time and the distance between receptors at which they are expressed in the membrane. Studying the distribution of ORs along the neurons will also contribute to understand the differences in OR performance (currently under investigation, Wicher, D., personal communication, December 2021).

## Materials and methods

### Fly rearing and fly lines

*Drosophila melanogaster* flies were reared under a 12 h light: 12 h dark cycle at 25° on conventional agar medium.

A list of all flies used can be found in **Table 1**.

**Table1.** List of flies lines used

|  | Genotype |
| --- | --- |
| 1 | *Canton-S* (WT) |
| 2 | *w;UAS-GCaMP6f;Or22a-Gal4* |
| 3 | *w;UAS-GCaMP6f;Or56a-Gal4/TM6B* |
| 4 | *CyO/BL;Or22a-Gal4, UAS-DenMark* |
| 5 | *CyO/BL;Or22a-Gal4, UAS-MitoGFP* |
| 6 | *(CyO)/+;Or56a-Gal4, UAS-DenMark* |
| 7 | *(CyO)/+;Or56a-Gal4, UAS-MitoGFP/TM6B* |

### Single sensillum recordings (SSR)

A set of ORs was chosen to account for the variability in the *Drosophila* OR repertoire. Neurons in ab3 and ab5 sensilla respond to food related odors and are broadly tuned, expect for ab5A which is narrowly tunned (Hallem and Carlson, 2006; Knaden et al., 2012). Ab4 sensillum class accounts for the aversive encoding neurons. Ab4A neuron is broadly tuned and Ab4B is very only tunned to one compound: geosmin. This specific compound has a great ecological value for the fly since its presence indicates harmful bacteria eliciting avoidance (Stensmyr et al., 2012). At1 houses the cVA sensing neuron, involved in social aggregation and sexual behaviors (Bartelt et al., 1985; Kurtovic et al., 2007). A summary of target sensilla and chemicals used for SSR can be found in **Table 2**.

**Table 2.** Summary of target sensillum and chemicals used for SSR experiments

| Sensillum type (neuron) | Compound | CAS |
| --- | --- | --- |
| ab3(A) | ethyl hexanoate | 123-66-0 |
| ab3(B) | 2-heptanone | 110-43-0 |
| ab4(A) | E2-hexanal | 6728-26-3 |
| ab4(B) | geosmin | 16423-19-1 |
| at1 | cVA | 6186-98-7 |
| ab5(A) | geranyl acetate | 105-87-3 |
| ab5(B) | pentyl acetate | 628-63-7 |

**Table 3.**
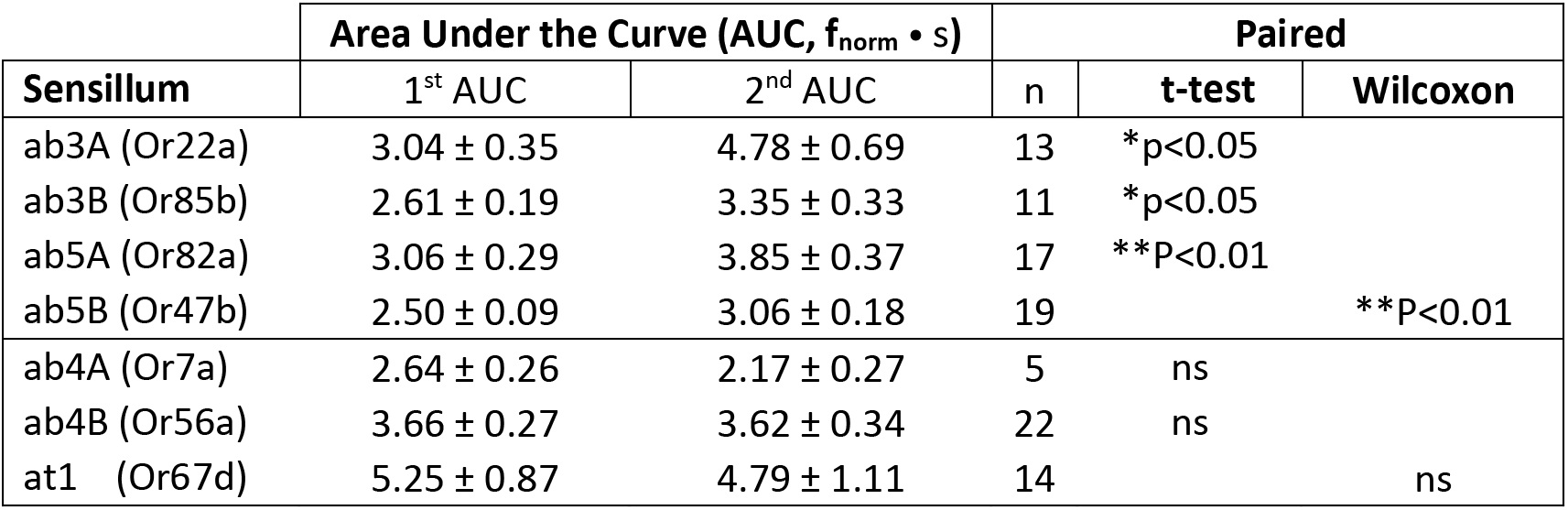
Sensitization results for SSR experiments. Significant difference between first and second AUC indicates sensitization. Data represent mean ± SEM.

**Table 4.** Results for response intensity (as the average increase in fluorescence ΔF/F_0_) after application of the OR ligands. Green indicates neuronal compartments of Or22a-expressing neurons, and red of Or56a-expressing neurons. Data is expressed as mean ± SEM

| Response intensity | Control |  | Auranofin |  |
| --- | --- | --- | --- | --- |
|  | 1<sup>st</sup> response | 2<sup>nd</sup> response | 1<sup>st</sup> response | 2<sup>nd</sup> response |
| Outer dendrites | 48.88 ± 20.38 | 71.34 ± 22.23 | 25.66 ± 9.87 | 9.55 ± 5.53 |
| Inner dendrites | 66.86 ± 13.91 | 103.8 ± 23.65 | 46.28 ± 9.18 | 15.99 ± 6.52 |
| Soma | 32.18 ± 8.18 | 50.47 ± 12.06 | 29.94 ± 5.55 | 21.17 ± 7.12 |
| Outer dendrites | 64.74 ± 15.36 | 34.89 ± 4.75 | 46.03 ± 11.3 | 14.15 ± 6.24 |
| Inner dendrites | 50.79 ± 14.12 | 35.79 ± 7.92 | 30.95 ± 10.72 | 21.95 ± 6.67 |
| Soma | 66.02 ± 16 | 45.14 ± 10.68 | 39.13 ± 7.5 | 23.13 ± 7.53 |

SSR experiments were carried out with wild-type *D. melanogaster* Canton-S (WTcs, stock #1), obtained from the Bloomington Drosophila Stock Center (www.flystocks.bio.indiana.edu). A 5-8-day old fly was held immobile in a 200 μl pipette tip and fixed on a glass side with laboratory wax. The funiculus (third antennal segment) was fixed in such position that either the medial or the posterior side faced the observer. Extracellular recordings were done using electrochemically (3M KOH) sharpened tungsten electrodes by inserting ground electrode in the eye and inserting recording electrode into the base of sensilla using micromanipulator system (Luigs and Nuemann SM-10). Sensilla were visualized with 1000x magnification using a binocular microscope (Olympus BX51WI). Signals were amplified (Syntech Uni-versal AC/DC Probe; www.syntech.nl), sampled (96000/s) and filtered (3kHz High-300Hz low, 50/60 Hz suppression) using a USB-IDAC. Neuronal activity was recorded using AutoSpike software (v3.7) for 3 seconds pre and 10 seconds post stimulus. Stimulus was delivered for 500 ms and was added to pre-humidified air being constantly delivered on the fly at a rate of 0.6 LPM.

Stimulus was prepared by pipetting 10 μl of the desired compound dissolved in hexane (or mineral oil for ab3 sensilla) onto a filter paper with a diameter of 10 mm and placed inside a glass Pasteur pipette. No more than 3 sensilla were recorded from each fly and odors were used once. Pared stimulations were 20 seconds apart, and there was 2-minute interval between pairs.

For odor application, a stimulus controller (Stimulus Controller CS-55, Syntech) was used, which produced a continuous airstream flow of 0.6 LPM air monitored by a flowmeter (Cole-Parmer, www.coleparmer.com). During stimulation, airflow bypassed a complementary air stream (0.6 l/min during 0.5 s) through the stimulus pipette placed roughly 3 cm from the preparation.

### Confocal imaging

By crossing fly lines 3*4 and 5*6 we generated flies with marked mitochondria (Mito-GFP in green) and a dendritic marker (DenMark (Nicolaï et al., 2010) in red) under the control of the OSN Or22a- or Or56a-Gal4 driver respectively. This allowed us to observe differences in mitochondrial distribution in the dendrites of this two different OSNs population.

Images were acquired on a cLSM 880 (Carl Zeiss, Oberkochen, Germany) using a 40x water immersion objective (C-Apochromat, NA: 1.2, Carl Zeiss) and adjusted for contrast and brightness by using LSM Image Browser 4.0 (Carl Zeiss).

### Antennal preparation

For calcium imaging experiments, antennae of 4-8 days old females were excised and prepared as described in Mukunda et al. 2014. Briefly, flies were anesthetized on ice, antennae were excised and fixed in vertical position with a two component silicone and finally immersed in *Drosophila* Ringer solution (in mM: HEPES, 5; NaCl, 130; KCl, 5; MgCl_2_, 2; CaCl_2_, 2; and sucrose, 36; pH = 7.3). After, funiculus was cut allowing access to the OSNs for experiments. Throughout the experiments, antennae were submerged in *Drosophila* Ringer solution.

### Calcium Imaging

A monochromator (Polychrome V, Till Photonics, Munich, Germany) coupled to an epifluorescence microscope (Axioskop FS, Zeiss, Jena, Germany) was employed for imaging. A water immersion objective (LUMPFL 60x/0.90; Olympus, Hamburg, Germany) controlled by an imaging control unit (ICU, Till Photonics) was used. Fluorescence images were acquired using a cooled CCD camera controlled by TILLVision 4.5 software (TILL Photonics).

GCaMP6f was exited with 475 nm light at 0.2 Hz frequency with an exposition time of 50 ms. Emitted light was separated by a 490 nm dichroic mirror and filtered with a 515 nm long-pass filter. TillVision software was used to subtract background fluorescence and to define regions of interest (ROI) characterized by a change in the [Ca^2+^]_i_ indicated by a change in fluorescence. The response magnitude was then calculated (ΔF/F_0_) in percentage following Mukunda et al. 2014.

Experiments lasted 15 min with a sampling interval of 5 seconds. Samples were continuously perfused during the experiments with bath solution in a perfusion/recording chamber (RC-27, Warner Instruments Inc., Hamden, CT, USA).

OSNs were stimulated by application of 10 μl of ethyl hexanoate (0.5 μM) or geosmin (1 nM) 2 cm away from the sample. Two control stimulations were performed 100 seconds apart. Then, solution was changed into *Drosophila* Ringer+Auranofin 25 μM and 2 other paired stimulations were performed.

### Chemicals

Auranofin (C_20_H_34_AuO_9_PS), ethyl hexanoate, (±)-Geosmin, 2-heptanone, E2-hexanal, geranyl acetate and pentyl acetate were purchased from Sigma Aldrich (Steinheim, Germany). 11-cis-Vaccenyl acetate (cVA) was purchased from Pherobank (Pherobank B.V., The Netherlands). All chemicals have ≥ 97% purity.

### Data analysis

SSR traces were analyzed using AutoSpike32 (Syntech NL 1998). For response kinetics (**Figure 1**), spike frequency ratios were analyzed as PSTH histograms in 25ms bins. By dividing each 25ms frequency by the average pre-stimulus frequency over 2 seconds, a normalized frequency ratio (f_norm_) per each time bin was obtained. PSTH represented in the figures show normalized means ± SEM for *n* paired stimulations. Between 2 and 4 flies were used for each odor, and no more than 3 sensilla per fly. To obtain response velocity the first derivative of f_norm_ was calculated. To test for the occurrence of sensitization the Area Under the Curve (AUC) was calculated for all sensilla. For comparison two-tailed paired t-test or Wilcoxon matched-pairs signed rank test was performed.

For fluorescence quantification of mitochondria (**Figure 3**) ImageJ was used following the method described by McCloy, R. A and colleagues (McCloy et al., 2014). Briefly, an outline was drawn around each soma and a transversal line through inner dendrites to measure area or width and mean fluorescence along with background. The corrected total cellular fluorescence (CTCF) = integrated density – (area of selected cell × mean fluorescence of background readings), was calculated. Two-tailed unpaired Student *t* tests were then performed.

Data analysis and graphs were generated using RStudio (Version 1.3.1093) and GraphPad Prism 9.

Figures were customized with Adobe Illustrator CS5 (Version 15).

## Authors contributions

DW and LH-L designed the experiments. LH-L and VPM conducted the experiments. LH-L performed the analysis. All authors discussed the results and wrote the article.

## Funding

This study was supported by the Max Planck Society (LH-L, VPM, BH and DW).

## Acknowledgments

We thank Dr. Vignesh Venkateswaran and Sinisa Prelic in generating the R code for calcium imaging analysis and assistance in data analysis.. We thank Dr. Ian Keesey for assistance and guidance at the initial steps of SSR experiments.

## Notes

### Competing Interest Statement

The authors have declared no competing interest.

