## Supplementary material for "Response plasticity of *Drosophila* olfactory sensory neurons"

Lorena Halty-deLeon, Venkatesh Pal Mahadevan, Bill S. Hansson‡ and Dieter Wicher‡*
Max Planck Institute for Chemical Ecology
Jena, Germany

‡Shared senior authorship

Supplementary figure 1

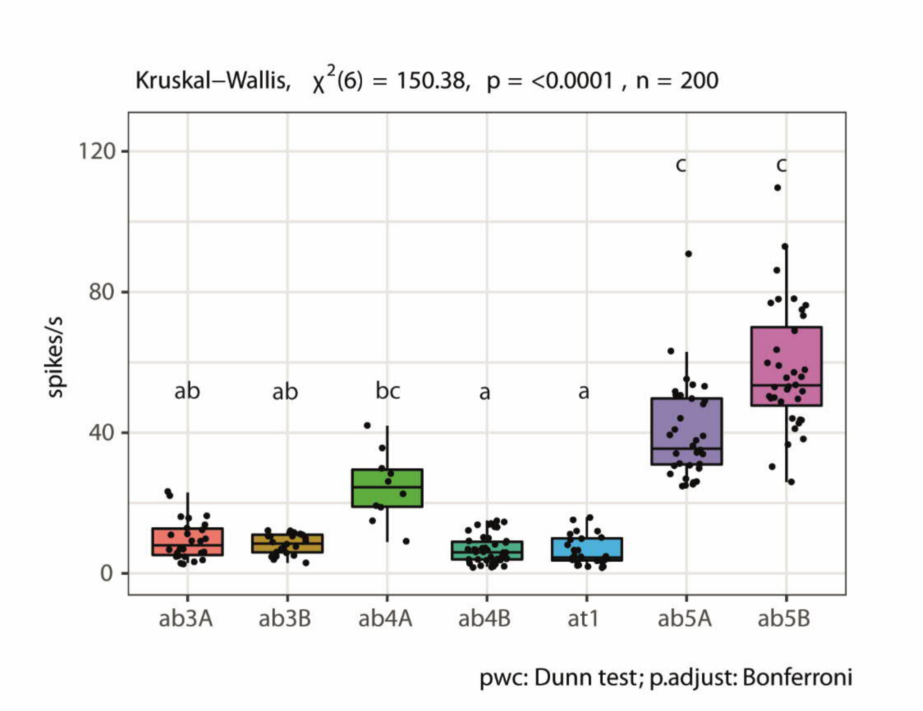


S1. Spontaneous activity. Box plots showing the spontaneous activity per second of the different neurons. Different small letters above the boxplots indicate significant differences (Kruskal-Wallis test followed by Dunn's pairwise (pwc) multiple comparison test. Data represent mean ± SEM. For detailed statistics, see Table S1.

Supplementary figure 2

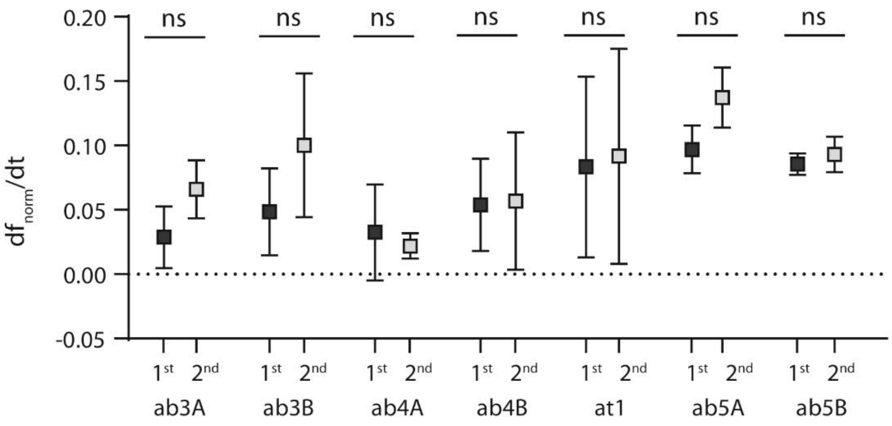


S2. Response velocity. Squares depict the maximum velocity of the first (black) and second (gray) response for each neuron. Data extracted as the first derivative ($df_{norm}/dt$) from curves in Figure 1B. Data represent maximum ± SEM, two tail t-test, ns not significant.

Supplementary figure 3

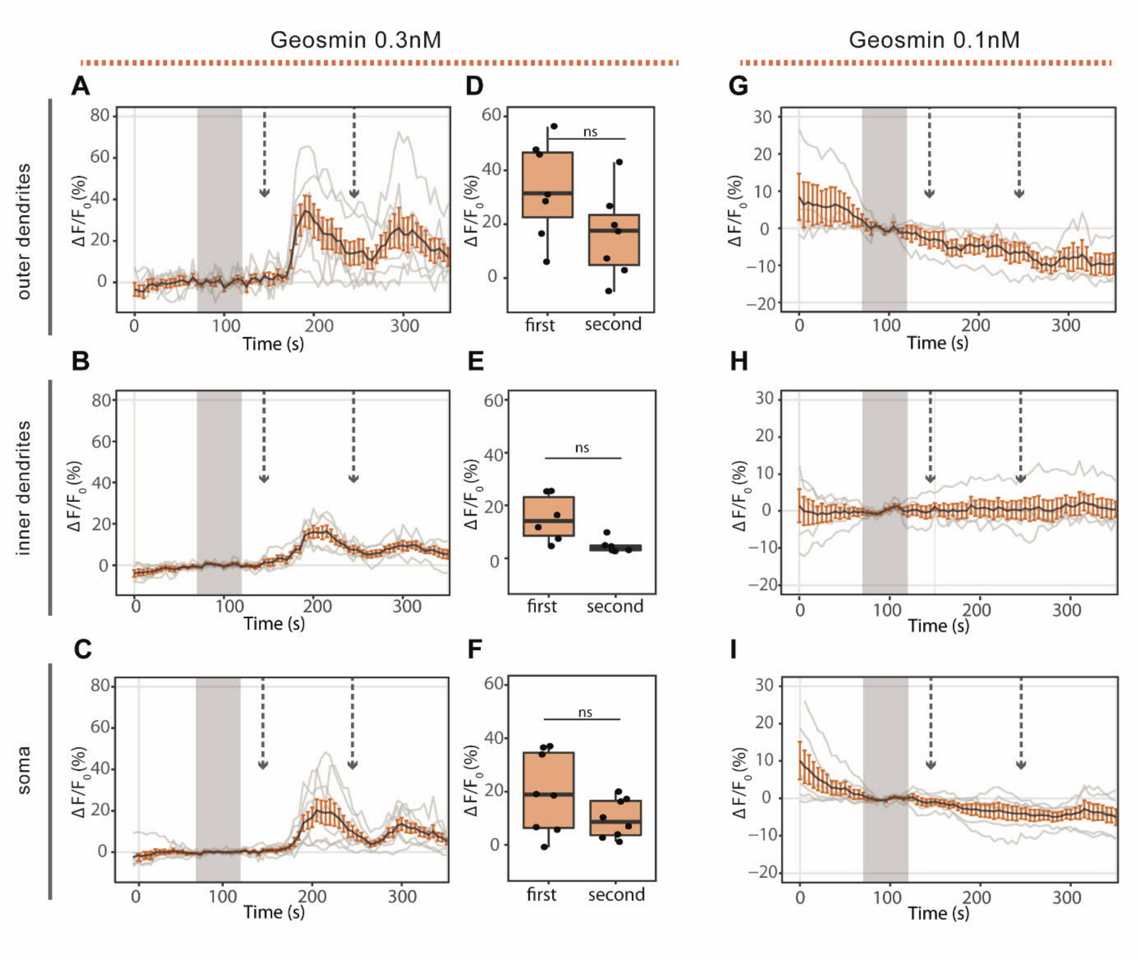


S2. No sensitization in Or56a neurons. A,B,C: Kinetics show averaged time course of the change in fluorescence intensity ($\Delta F/F_0$) in *Drosophila* OSNs after application of 0.3 nM geosmin (arrows). At 0.3 nM there is no difference between applications in outer dendrites (A, n=7), inner dendrites (B, n=6) and soma (C, n=8). D,E,F: maximum increase in $\Delta F/F_0$ after geosmin application in the different compartments as in A-C. G,H,I: No responses are observed after 0.1 nM geosmin application (arrows). Gray bar indicates where data was normalized to obtain $\Delta F/F_0$. Data represent mean ± SEM; two-tail paired t-test, ns not significant.

Supplementary table 1

Table S1 Results from Dunn´s multiple comparison test with Bonferroni correction following a Kruskal-Wallis test. Spikes/s show mean + SEM

| group1 | group2 | $n_1$ | Spikes/s | $n_2$ | Spikes/s | statistic | p.adj | p.adj.signif |
|---|---|---|---|---|---|---|---|---|
| ab3A | ab3B | 26 | 9.58 ± 1 | 22 | 8.32 ± 0.6 | -0.11052 | 1 | ns |
| ab3A | ab4A | 26 | 9.58 ± 1 | 10 | 24.70 ± 3.1 | 2.472702 | 0.281601 | ns |
| ab3A | ab4B | 26 | 9.58 ± 1 | 44 | 6.84 ± 0.6 | -1.29403 | 1 | ns |
| ab3A | ab5A | 26 | 9.58 ± 1 | 34 | 40.06 ± 2.38 | 5.245598 | 3.27E-06 | **** |
| ab3A | ab5B | 26 | 9.58 ± 1 | 36 | 57.95 ± 2.9 | 6.853514 | 1.51E-10 | **** |
| ab3A | at1 | 26 | 9.58 ± 1 | 28 | 6.32 ± 078 | -1.52926 | 1 | ns |
| ab3B | ab4A | 22 | 8.32 ± 0.6 | 10 | 24.70 ± 3.1 | 2.496476 | 0.263412 | ns |
| ab3B | ab4B | 22 | 8.32 ± 0.6 | 44 | 6.84 ± 0.6 | -1.10326 | 1 | ns |
| ab3B | ab5A | 22 | 8.32 ± 0.6 | 34 | 40.06 ± 2.38 | 5.111615 | 6.71E-06 | **** |
| ab3B | ab5B | 22 | 8.32 ± 0.6 | 36 | 57.95 ± 2.9 | 6.636389 | 6.75E-10 | **** |
| ab3B | at1 | 22 | 8.32 ± 0.6 | 28 | 6.32 ± 078 | -1.34952 | 1 | ns |
| ab4A | ab4B | 10 | 24.70 ± 3.1 | 44 | 6.84 ± 0.6 | -3.54014 | 0.008398 | ** |
| ab4A | ab5A | 10 | 24.70 ± 3.1 | 34 | 40.06 ± 2.38 | 1.241198 | 1 | ns |
| ab4A | ab5B | 10 | 24.70 ± 3.1 | 36 | 57.95 ± 2.9 | 2.360504 | 0.383253 | ns |
| ab4A | at1 | 10 | 24.70 ± 3.1 | 28 | 6.32 ± 078 | -3.62818 | 0.005994 | ** |
| ab4B | ab5A | 44 | 6.84 ± 0.6 | 34 | 40.06 ± 2.38 | 7.386824 | 3.16E-12 | **** |
| ab4B | ab5B | 44 | 6.84 ± 0.6 | 36 | 57.95 ± 2.9 | 9.273146 | 3.80E-19 | **** |
| ab4B | at1 | 44 | 6.84 ± 0.6 | 28 | 6.32 ± 078 | -0.39877 | 1 | ns |
| ab5A | ab5B | 34 | 40.06 ± 2.38 | 36 | 57.95 ± 2.9 | 1.66126 | 1 | ns |
| ab5A | at1 | 34 | 40.06 ± 2.38 | 28 | 6.32 ± 078 | -6.98715 | 5.89E-11 | **** |
| ab5B | at1 | 36 | 57.95 ± 2.9 | 28 | 6.32 ± 078 | -8.65314 | 1.05E-16 | **** |
